# Cold-hearted bats: Cardiac function and metabolism of small bats during torpor at subzero temperatures

**DOI:** 10.1101/192526

**Authors:** Shannon E. Currie, Clare Stawski, Fritz Geiser

**Affiliations:** Centre for Behavioural and Physiological Ecology, Zoology, University of New England, Armidale, 2351, NSW, Australia; Leibniz Institute for Zoo and Wildlife Research, Alfred-Kowalke-Str. 17, 10315 Berlin, Germany

**Author notes:** **Corresponding Author**: Shannon. E. Currie, Leibniz Institue for Zoo and Wildlife Research, Alfred-Kowalke-Str. 17, 10315 Berlin, Germany, **Email**.

**Keywords:** bats, heart rate, hibernation, metabolism, oxygen pulse, thermoregulation, torpor

## Abstract

Despite their small size and large relative surface area, many hibernating bats have the ability to thermoregulate and defend their body temperature (T_b_) often below 10°C by an increase in metabolic rate. Above a critical temperature (T_crit_) animals usually thermoconform. We investigated the physiological responses above and below T_crit_ for a small tree dwelling bat (*Chalinolobus gouldii*, ∼14g) that is often exposed to subzero temperatures during winter. Through simultaneous measurement of heart rate (HR) and oxygen consumption (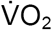) we show that the relationship between oxygen transport and cardiac function is substantially altered in thermoregulating torpid bats down to −2°C, compared with thermoconforming torpid bats at mild ambient temperatures (T_a_ 5-20°C). T_crit_ for this species was T_a_ 0.7 ± 0.4°C, with a corresponding T_b_ of 1.8 ± 1.2°C. Below this T_crit_ animals began to thermoregulate, indicated by a considerable but disproportionate increase in both HR and 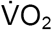. The maximum increase in HR was only 4-fold greater than the average thermoconforming minimum, compared to a 46-fold increase in 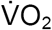. The differential response of HR and 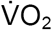 to low T_a_ was represented by a 15-fold increase in oxygen delivery per heart beat (cardiac oxygen pulse). During torpor at low T_a_, thermoregulating bats maintain a relatively slow HR and compensate for increased metabolic demands by significantly increasing stroke volume and tissue oxygen extraction. Our study provides valuable new information on the relationship between metabolism and HR in an unstudied physiological state and further advances our knowledge of the thermogenic capacity of small bats.

## Introduction

Extremely low ambient temperatures (T_a_) create an energetic hurdle for many animals, particularly small endotherms that must markedly increase their heat production to maintain a constant high body temperature (T_b_). Torpor, a controlled reduction of metabolic rate (MR), heart rate (HR) and T_b_, is therefore critical for many small mammals to save energy during inclement conditions (Ruf & Geiser 2015). During torpor the hypothalamic control for temperature regulation is adjusted to a new, lower minimum T_b_ that is regulated at or just above a critical external temperature (T_crit_). Depending on ambient conditions, animals will thermoconform down to this T_b_/T_a_ (Heller 1979; Geiser 2011). However, when T_a_ falls below T_crit_ animals can either rewarm entirely to a normothermic T_b_ or remain torpid and increase metabolic heat production to thermoregulate and defend their minimum T_b_.

Details regarding thermoregulation during torpor at subzero temperatures are generally scant and are most often restricted to medium-sized northern or arctic species such as ground squirrels (Geiser & Kenagy 1988; Buck & Barnes 2000; Richter *et al.* 2015). However, small hibernators, such as insectivorous bats often use torpor year round in temperate climates where they are exposed to subzero temperatures during winter (Eisentraut 1956; Davis & Reite 1967; Masing & Lutsar 2007; Wojciechowski, Jefimow & Tegowska 2007). Thermoregulation during torpor, while still conserving substantial amounts of energy, can be expensive when used over long periods and against a large T_b_-T_a_ differential (Karpovich, Tøien, Buck & Barnes 2009; Richter *et al.* 2015). At extremely low T_a_, defending T_b_ above T_crit_ likely requires an alteration of a number of physiological components essential to thermoregulation. While studies have been conducted on thermal energetics of torpor at low Ta, detailed information on cardiac function and its relationship to metabolism are entirely lacking.

The cardiovascular system is vital for maintaining physiological processes during torpor as the heart regulates the supply of blood gases, nutrients and hormones to the body. During torpor at low T_b_ the heart must work to facilitate adequate blood supply under conditions of reduced flow and slow ventilation/low oxygen intake (Milsom, Zimmer & Harris 2001). In addition, in order to minimise metabolic costs during torpor, adequate perfusion is only retained in vital tissues and organs due to circulatory adjustments (Carey, Andrews & Martin 2003). The hearts of hibernating animals are adapted to function at low T_b_ and are capable of withstanding temperatures well below the critical point for fibrillation and death in non-hibernating species (Lyman & Blinks 1959; van Veen, van der Heyden & van Rijen 2008). It is widely known that during torpor in thermoconforming hibernators the heart continues to beat in a co-ordinated rhythm and is resistant to detrimental arrhythmia (Johansson 1996; van der Heyden & Opthof 2005). However, when animals are forced to thermoregulate during torpor the cardiovascular system must be altered alongside increases in metabolism.

We investigated the thermal physiology, in particular interrelations between HR and MR below and above the T_crit_, for torpid *Chalinolobus gouldii*, a common tree-dwelling bat species whose distribution extends across Australia (Churchill 2008). While temperate zone northern hemisphere bats tend to hibernate in thermally stable caves during winter, many southern hemisphere bats often hibernate in tree hollows or under bark. These roosts are thermally labile and often experience subzero T_a_ (Law & Chidel 2007; Turbill 2008; Clement & Castleberry 2012; Stawski & Currie 2016). Although there are some data on thermal biology of *C. gouldii* (Hosken & Withers 1997; Codd *et al.* 2000) information concerning cardiac function and its relationship to metabolism and thermal energetics in this species, or at such low temperatures in any bat, is entirely lacking. Moreover, explicit investigations of T_crit_ and thermogenic capacity in hibernating bats are extremely limited (Reite & Davis 1966; Stawski & Geiser 2011).

As *C. gouldii* regularly experience low T_a_, we hypothesised that the T_crit_ for this species would be close to 0°C and that HR would be substantially reduced during steady-state torpor, with corresponding low T_b_ and MR, until T_a_ reached T_crit_. However, below T_crit_ we hypothesised that bats would increase both HR and MR, but maintain a steady T_b_.

## Materials and methods

Six non-reproductive *Chalinolobus gouldii* (3 females and 3 males; capture weight 14.3 ± 2.6g) were collected from a residential roof near the University of New England (UNE) in Armidale, NSW, during the austral autumn (April) 2015. Individuals were kept in outdoor aviaries at UNE for three months and measurements were conducted from May - July 2015 (austral late Autumn/Winter). Food *(Tenebrio molitor* larvae dusted with a supplement of Wombaroo(™) Insectivore Mix) and water were provided *ad libitum.* Prior to release at their capture site, animals were fitted with temperature sensitive radio-transmitters as part of another study (Stawski & Currie 2016).

### Respirometry and electrocardiogram measurements

In the late afternoon, bats were placed in air tight respirometry chambers in a temperature controlled cabinet where they remained overnight and into the following day(s). Animals were not given access to food or water during respirometry measurements to ensure they were post absorptive. Bats were weighed to the nearest 0.1g before and after measurements. The T_a_ in the chamber was recorded to the nearest 0.1°C using a calibrated thermocouple placed 5mm within the chamber and read using a digital thermometer.

Respirometry chambers (0.53 litres) were made from modified polycarbonate enclosures with clear lids, lined with a small patch of hessian cloth (burlap) from which the bats could roost. Air flow through the chambers was controlled by rotameters and measured with mass flowmeters (Omega FMA-5606; Stamford, CT, USA) that were calibrated prior to the start of experimentation. Flow rate (200 ml/min) was adjusted to ensure that 99% equilibrium was reached within 12 min. Oxygen concentration was measured using a FC-1B Oxygen Analyser (Sable Systems International Inc., Las Vegas, USA, resolution 0.001%). Measurements were taken from the chamber every minute for 15 min and then switched to outside air for reference readings (3 min) using solenoid valves. Digital outputs of the oxygen analyser, flowmeter and digital thermometer were recorded on a PC using custom-written data-acquisition software (G. Körtner). 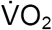 values were calculated per minute using standardised gas volumes and equation 3a of Withers (1977) assuming a respiratory quotient of 0.85.

Electrocardiograms (ECG) were recorded from two leads attached to adhesive electrodes on the forearms of each bat following the methods of Currie, Körtner and Geiser (2014). Attachment took place once bats were torpid, either following lights on in the morning or during the night following rectal T_b_ measurements. All bats returned to torpor within 1h of lead attachment. Electrocardiograms were recorded using LabChart v7.3 software and analysed to determine HR by calculating instantaneous HR per second. HR data was then exported to excel for further analysis.

### T_a_ above zero

For measurements of steady-state thermoconforming torpor at mild T_a_ bats were placed in respirometry chambers overnight at T_a_ between 5 and 15°C. Following ECG lead attachment, and a return to steady-state torpor, the T_a_ of the chamber was increased by 2°C and once the new T_a_ was reached, this was left unchanged for at least 2h. The T_a_ was then progressively increased until animals rewarmed, either in response to changing T_a_ or the stimulus of lights off in the evening.

### T_a_ below zero

Bats were placed in respirometry chambers overnight at a T_a_ between 1 and 5°C. Following lights on and/or the attachment of ECG leads, T_a_ was progressively decreased in 1°C increments below 3°C. Individuals were exposed to each temperature for a minimum of 2h before the temperature was further reduced. Previous investigators (Reite & Davis 1966;S.E. Currie personal observ.) have shown that swift reductions in T_a_ (≥3-5°C) often induce arousal from hibernation in a number of species. Therefore care was taken to ensure T_a_ was gradually reduced to obtain an accurate representation of thermoregulatory T_crit_ and minimum T_a_ prior to spontaneous arousal. The minimum temperature reached before animals regularly aroused was −2°C, and therefore no measurements took place below this temperature.

### T_b_ measurements

T_b_ was measured to the nearest 0.1°C using a digital thermometer (Omega, HH-71T) and calibrated thermocouple probe inserted ∼2cm into the rectum. Rectal T_b_ was recorded for animals that had been torpid for at least 2h, indicated by a low MR and HR (n=6, N=26). Measurements were made within 1 min of removing the bat from the respirometry chamber and since the rate of rewarming from torpor is initially slow, rectal T_b_ was considered to be within 0.5°C of Tb prior to the disturbance.

### Analysis and statistics

Bats were considered torpid when 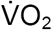 fell to less than 75% of the RMR at the same T_a_. During steady-state torpor HR and 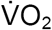 were averaged over the same time period, for at least 30 min,corresponding to minimum torpid 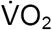. Bats were deemed to be thermoconforming when T_b_ was within 2°C of T_a_. When T_b_ was not available, individuals were considered to be thermoconforming when 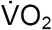 values fell to that, or below that, of bats with a T_b_ within 2°C of T_a_, at the same T_a_. The Q_10_ for rates of 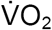 or HR of thermoconforming torpid bats was calculated using the following equation;

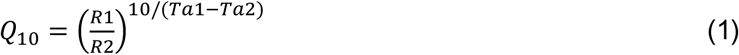

where R is the 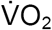 or HR at a particular T_a_ (1 or 2). Animals were considered to be thermoregulating in torpor when minimum 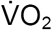 was at least double that of minimum average thermoconforming values at the same T_a_. Oxygen pulse (OP) was calculated for torpid bats by dividing the average 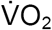 (ml/min) by corresponding average HR (bpm). The percentage contribution of HR to increases in oxygen transport in thermoregulating bats was calculated using the equation of Bartholomew and Tucker (1963);

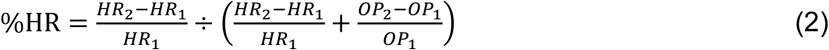

using HR and OP at T_a_ below 1°C (T_a_ 1 = 0.5°C; T_a_ 2 = −2.1°C).

Average resting HR (RHR) and 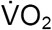 were calculated, from the period following arousal, for at least 5 min when 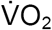 had fallen to ≤75% of maximum 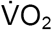 at the peak of rewarming. Unfortunately, during arousal at low T_a_ shivering associated with rewarming caused considerable artefact on ECG recordings, making calculation of HR extremely difficult. In addition, animals often moved during the final stages of rewarming, which resulted in detachment of ECG electrodes and cessation of HR recording. Therefore, data for RHR were only available for three bats across five T_a_.

All statistical analyses were performed using R v3.1.3 (R Core Team 2014), with a significance level of p<0.05. Means are presented ± standard deviations for number of animals (n) and number of observations (N). An analysis of covariance (ANCOVA) was undertaken to assess whether minimum 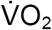 was significantly different following disturbance associated with ECG lead attachment. Linear mixed effects models (nlme package; Pinheiro *et al.* 2014) were used to assess the relationship between HR or 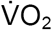 and T_a_ in thermoregulating torpid bats and resting bats. In order to determine any differences in slope between regressions for resting and thermoregulating 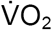 against T_a_, an ANCOVA was performed using nlme with individuals as a random factor. For comparisons between the slopes of HR and 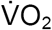 data were log transformed prior to ANCOVA using nlme, again with individual as a random factor. Standardised major axis regressions were performed (smatr package; Warton, Duursma, Falster & Taskinen 2012) to assess the correlation between 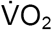 and HR when animals were either thermoconforming or thermoregulating during torpor. Pseudo-replication was accounted for by using the degrees of freedom as for mixed effect linear modelling adjusted for repeated measures. An ANCOVA using smatr was completed to determine whether there was a significant difference in the slopes for HR against 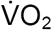 between the two torpid states.

All procedures were approved by the University of New England Animal Ethics Committee and New South Wales National Parks and Wildlife Service.

## Results

### Rest

At rest, following arousal from torpor, HR and 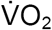 of normothermic bats increased with decreasing T_a_ in a qualitatively similar linear pattern (Figs 1A & B). Resting 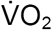 ranged from 1.44 to 10.04 ml g^−1^h^−1^ as T_a_ fell from 24.1 to −0.4°C (Fig 1A). Over a similar T_a_ range (19.4- 1.7°C), RHR increased from 412 to 698 bpm (Fig 1B), however the relationship between HR and T_a_ was not statistically significant (nlme; r^2^ = 0.96, p = 0.07).

**Figure 1.**
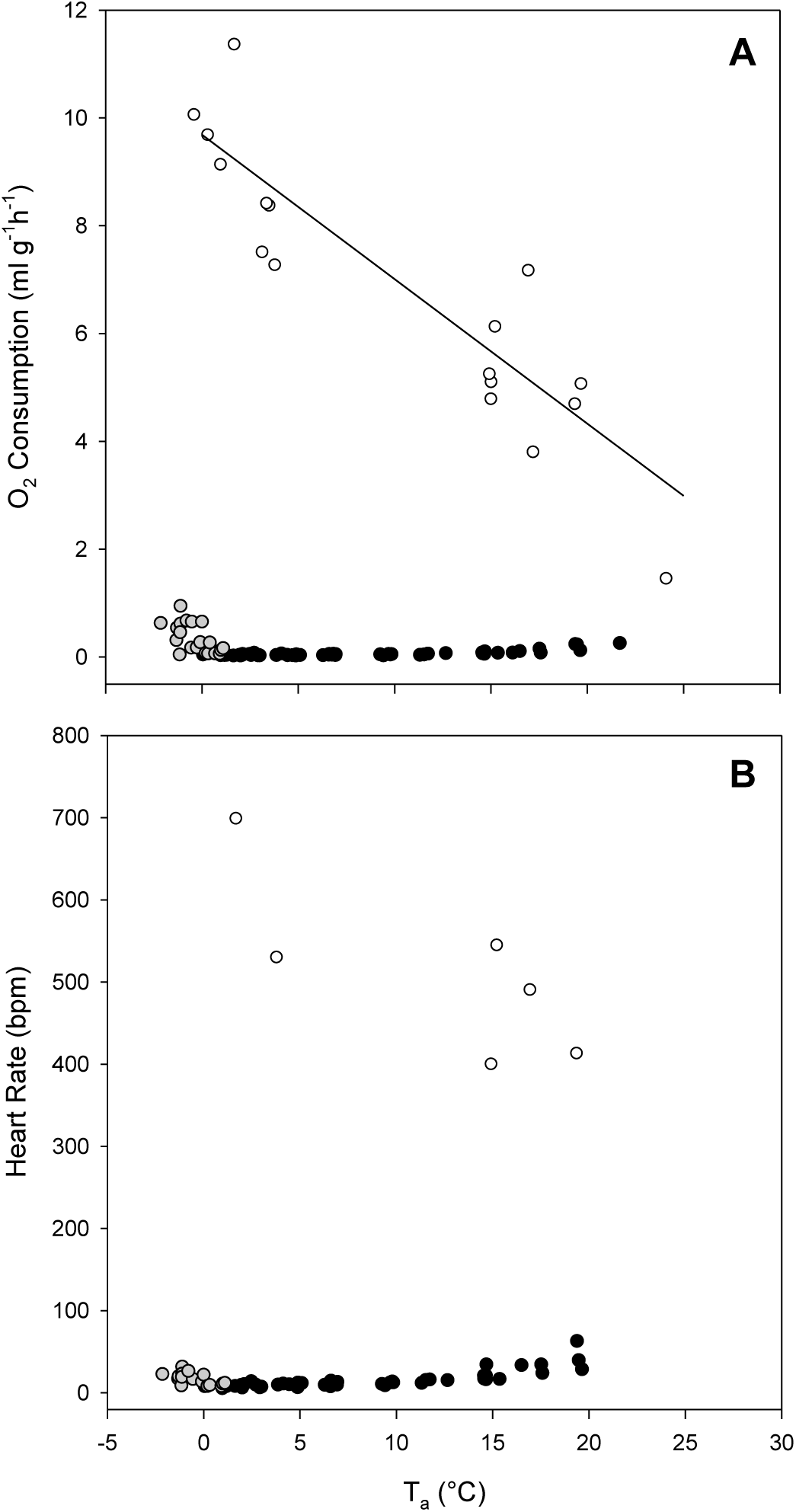
Oxygen consumption (A) and heart rate (B) as a function of ambient temperature (T_a_) for normothermic *C. gouldii* at rest (white circles, solid line; 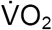 = 9.69 - 0.27(T_a_), r^2^=0.91, p<0.001), and during thermoconforming torpor (black circles) or thermoregulating torpor (grey circles).

### Thermoconforming torpid bats

All bats entered torpor at all T_a_ tested and reached low values indicative of steady-state torpor within 3h of disturbance associated with ECG lead attachment. There was no noteworthy impact of this disturbance as minimum 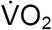 following ECG lead attachment was not significantly different from the minimum 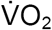 prior (p=0.85). Below T_a_ of 20°C most individuals remained torpid until the stimulus of lights off, except for a few bats that hibernated for up to 48h, or rewarmed in response to very low T_a_. Importantly, all bats were capable of spontaneously rewarming from the lowest T_a_ they were exposed to in our study.

When bats were in steady-state torpor and thermoconforming, 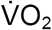 and HR followed T_a_ in a curvilinear manner. T_b_ fell to within 1.0 ± 0.9°C of T_a_ in thermoconforming individuals (n=6, N=20).The minimum average 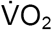 of thermoconforming bats was 0.02 ± 0.01 ml g^−1^ h^−1^ (n=6, N=9) recorded at 2.1 ± 0.3°C, which was <1% of resting values at a similar T_a_ (1.4 ± 0.5°C, V0_2_=10.23 ± 1.58 ml g^−1^ h^−1^, n=2, N=2). HR fell to an absolute minimum of 3bpm at T_a_ = 1.0°C while the average minimum torpor HR (THR) was 8 ± 2 bpm (n=6, N=7) and only 1.1% of RHR recorded at 1.4°C (698bpm, n=1, N=1). Even when bats were torpid at mild T_a_ (19.6°C) HR and 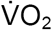 were <11% of the corresponding resting rates (THR = 10.3%; 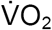 = 3.7%). When 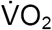 was plotted against HR in thermoconforming individuals there was a strong linear correlation (r^2^=0.88, p<0.001; Fig 2). On occasion, animals maintained a low 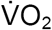 and HR when T_a_ fell below 1°C and one animal thermoconformed down to a T_a_ of −1 °C with a T_b_ of 0.6°C and corresponding 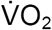 equal to 0.03ml g^−1^h^−1^ and HR of 7bpm.

**Figure 2.**
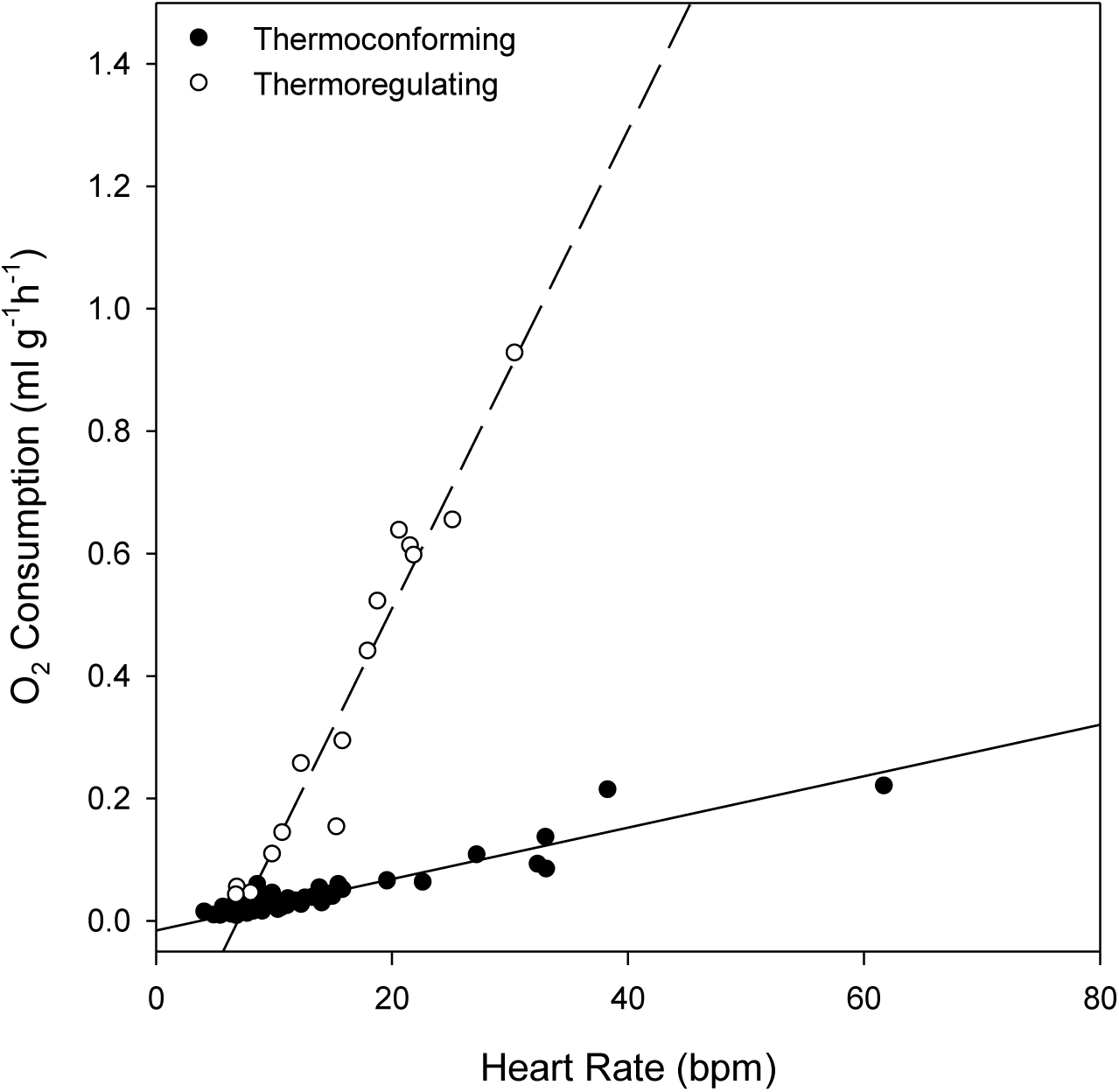
The relationship between heart rate (HR) and oxygen consumption in torpid *C. gouldii* for thermoconforming individuals (black circles, solid line; 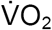=0.004(HR) - 0.016, r^2^=0.88, p<0.001) or thermoregulating individuals (white circles, dashed line; 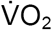=0.039(HR) - 0.272, r^2^=0.95, p<0.001).

The transport of oxygen per heart beat, or oxygen pulse showed a similar qualitative pattern to HR and 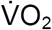 in bats during steady-state torpor, however declined only slightly with decreasing T_a_ down to 2°C (n=6, N=53; Fig 3). The Q_10_ for 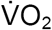 was 3.8 in thermoconforming bats, determined between T_a_ of 19.6 and 2.1°C. When compared to the basal metabolic rate (BMR) previously determined for this species (1.44 ml g-1 h-1; Hosken & Withers 1997) average Q_10_ was also 3.8. However, the corresponding Q_10_ for HR during thermoconforming torpor was only 2.6.

**Figure 3.**
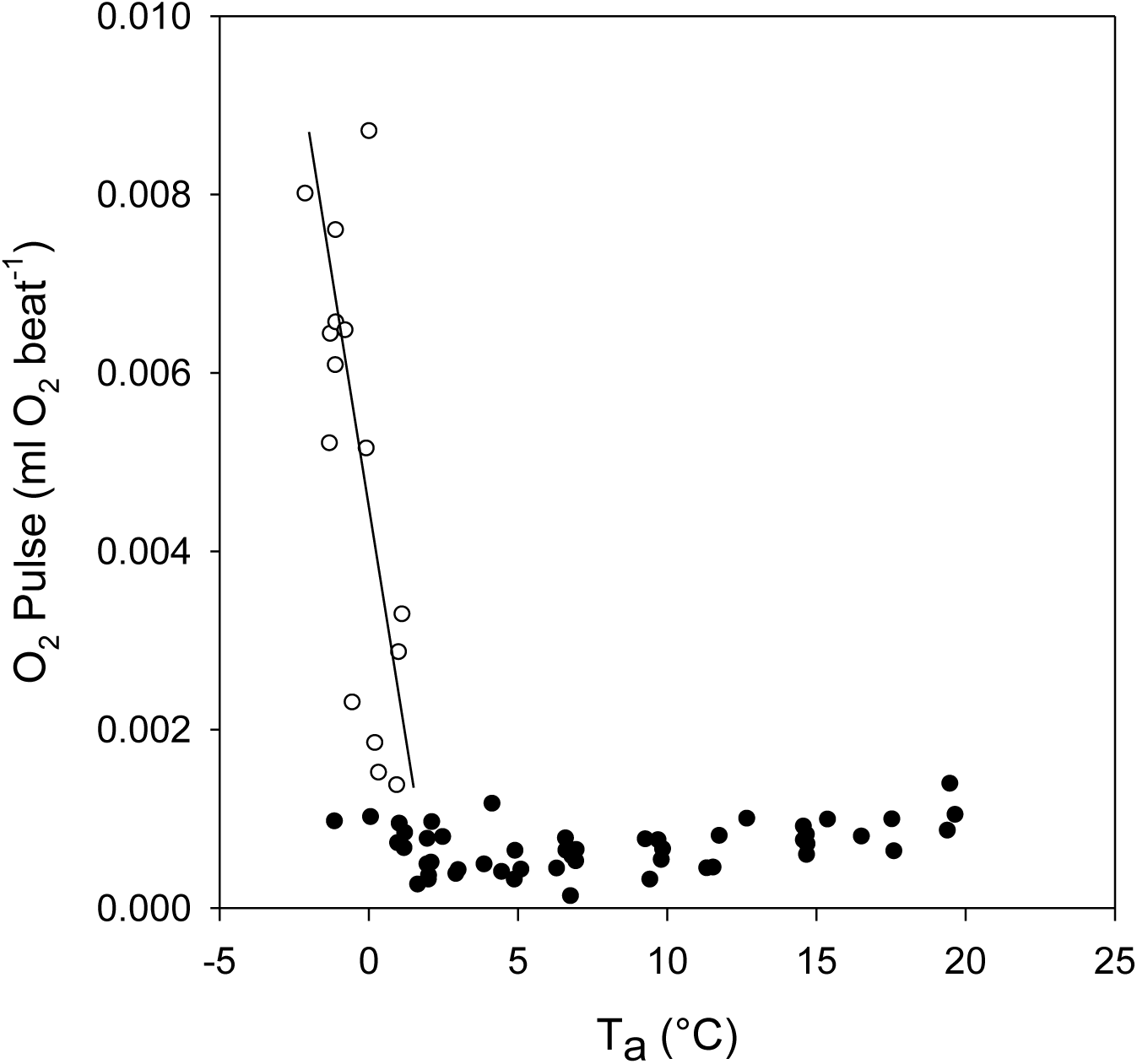
Oxygen pulse (O_2_P) as a function of ambient temperature (T_a_) in *C. gouldii* during thermoconforming torpor (black circles) or thermoregulating torpor (white circles). The relationship with Ta changes from curvilinear to linear when animals begin thermoregulating (solid line; O_2_P = 0.0045-0.0021 (Ta), r^2^ = 0.87, p<0.001)

### Thermoregulating torpid bats

Below a T_crit_ of 0.7 ± 0.4°C (n=6, N=7) both HR and 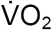 showed a substantial increase and bats began thermoregulating (Figs 1A & B). Average T_b_ at T_crit_ was 1.8 ± 1.2°C (n=5, N=5), and as T_a_ fell to −2°C, bats defended T_b_ at 2.3 ± 1.6°C (n=2, N=2). The majority of animals rewarmed spontaneously when T_a_ fell below −1.3°C, however one animal remained torpid down to a T_a_ of −2.1°C (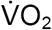 = 0.61 ml g^−1^h^−1^; HR = 22bpm). Thermoregulating torpid bats exposed to T_a_s below zero showed a steep increase in 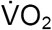 up to 46-fold when compared to minimum 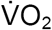 during torpor at T_a_ 2°C (n=6, N=15). In contrast, HR in these torpid thermoregulating bats only increased on average 2-fold (range 1 to 4-fold) over the same T_a_ range (n=6, N=15). The maximum HR recorded in thermoregulating torpid bats was 31bpm at T_a_ −1.1°C with a corresponding 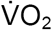 of 0.93 ml g^−1^ h^−1^.

There was a strong linear correlation between HR and 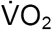 in thermoregulating bats (r^2^ = 0.95, p<0.001) and this was significantly steeper than the relationship for thermoconforming bats at T_a_ greater than 1°C (r^2^ = 0.88, p<0.001; ANCOVA, p<0.001; Fig 2).

Both HR and 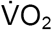 showed a significant linear increase with decreasing T_a_ in thermoregulating individuals (HR, r^2^ = 0.71, p<0.01; MR, r^2^ = 0.61, p<0.01; Figs 4 & 5). However, following log transformation, 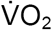 showed a significantly steeper response to declining T_a_ than HR (ANCOVA; p<0.05; Fig 6). This disproportionate increase in HR and 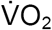 was also expressed via a significant linear increase in oxygen pulse as T_a_ fell below ∼1°C (r^2^=0.89, p<0.001; Fig 3). There was an almost 15-fold increase in average oxygen pulse from 2°C to −2°C (5.5 × 10^−4^ ml O_2_ beat^−1^ to 80 × 10^−4^ ml O_2_ beat^−1^ respectively) and the contribution of HR to changes in oxygen transport was minimal, at only 31% (calculated using Eq 2). Interestingly, the slope of the relationship between 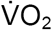 and T_a_, an indicator of thermal conductance, did not differ during thermoregulating torpor from that of resting bats (ANCOVA; p=0.63; Fig 4).

**Figure 4.**
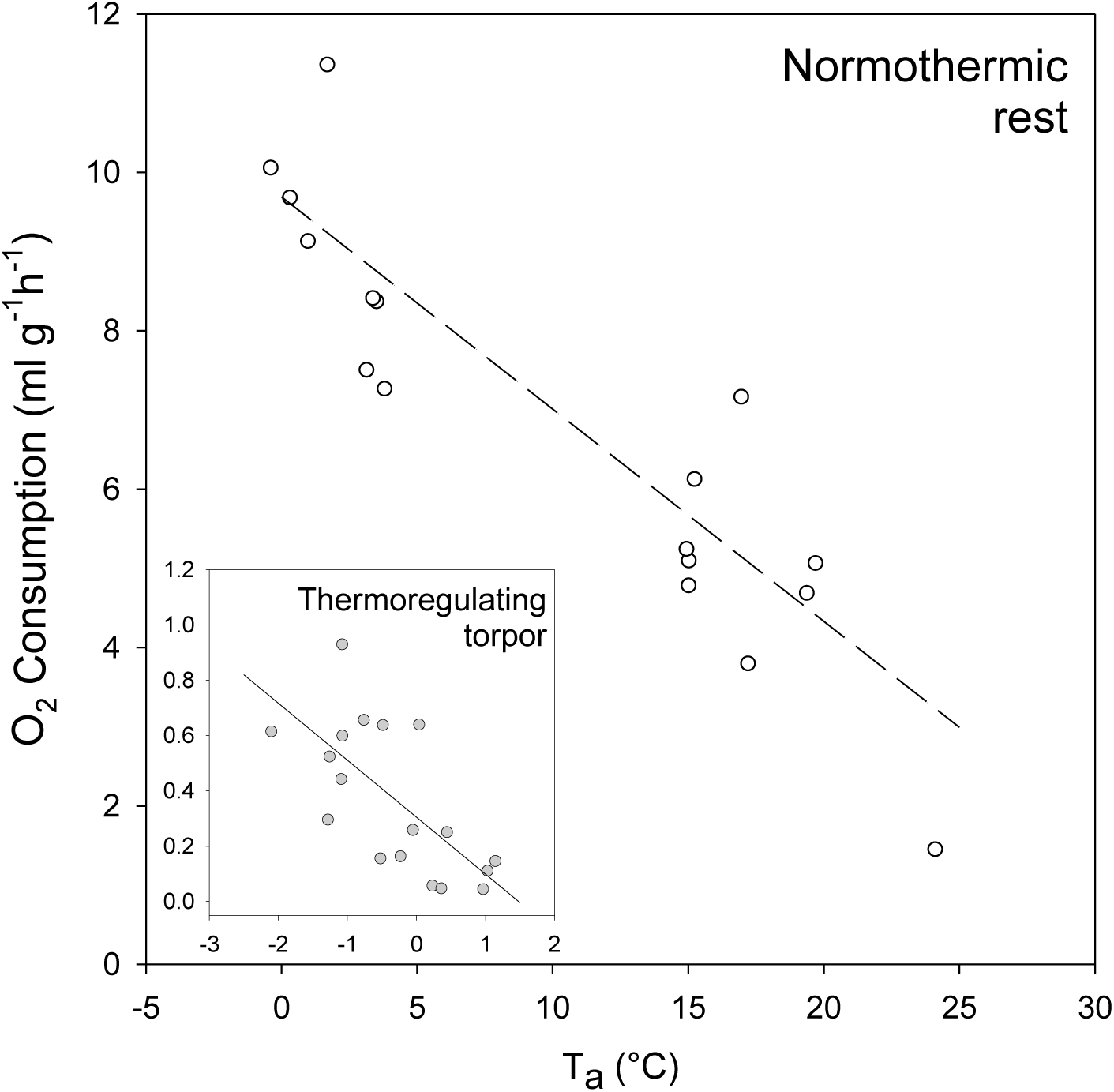
Oxygen consumption as a function of ambient temperature (T_a_) in thermoregulating *C. gouldii* at rest (white circles, dashed line; 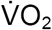 = 9.69 - 0.27(T_a_), r^2^=0.91, p<0.001), or during torpor (inlaid plot: grey circles, solid line; 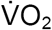 = 0.31 - 0.21(T_a_), r^2^=0.62, p<0.01).

**Figure 5.**
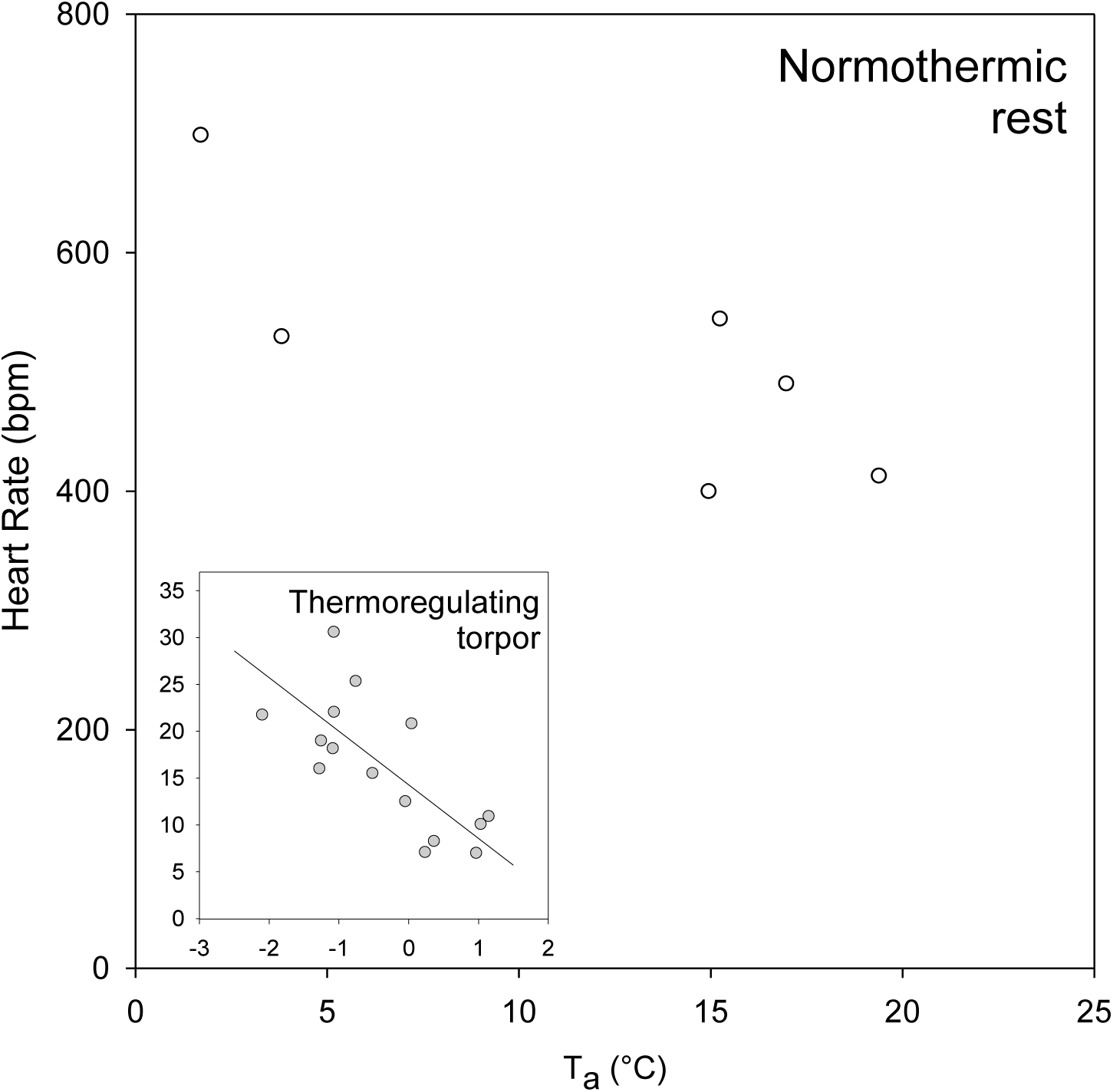
Heart rate as a function of ambient temperature (T_a_) in thermoregulating *C. gouldii* at rest (white circles), or during torpor (inlaid plot: grey circles, solid line; HR = 14.27- 5.72(Ta), r^2^=0.71, p<0.01).

**Figure 6.**
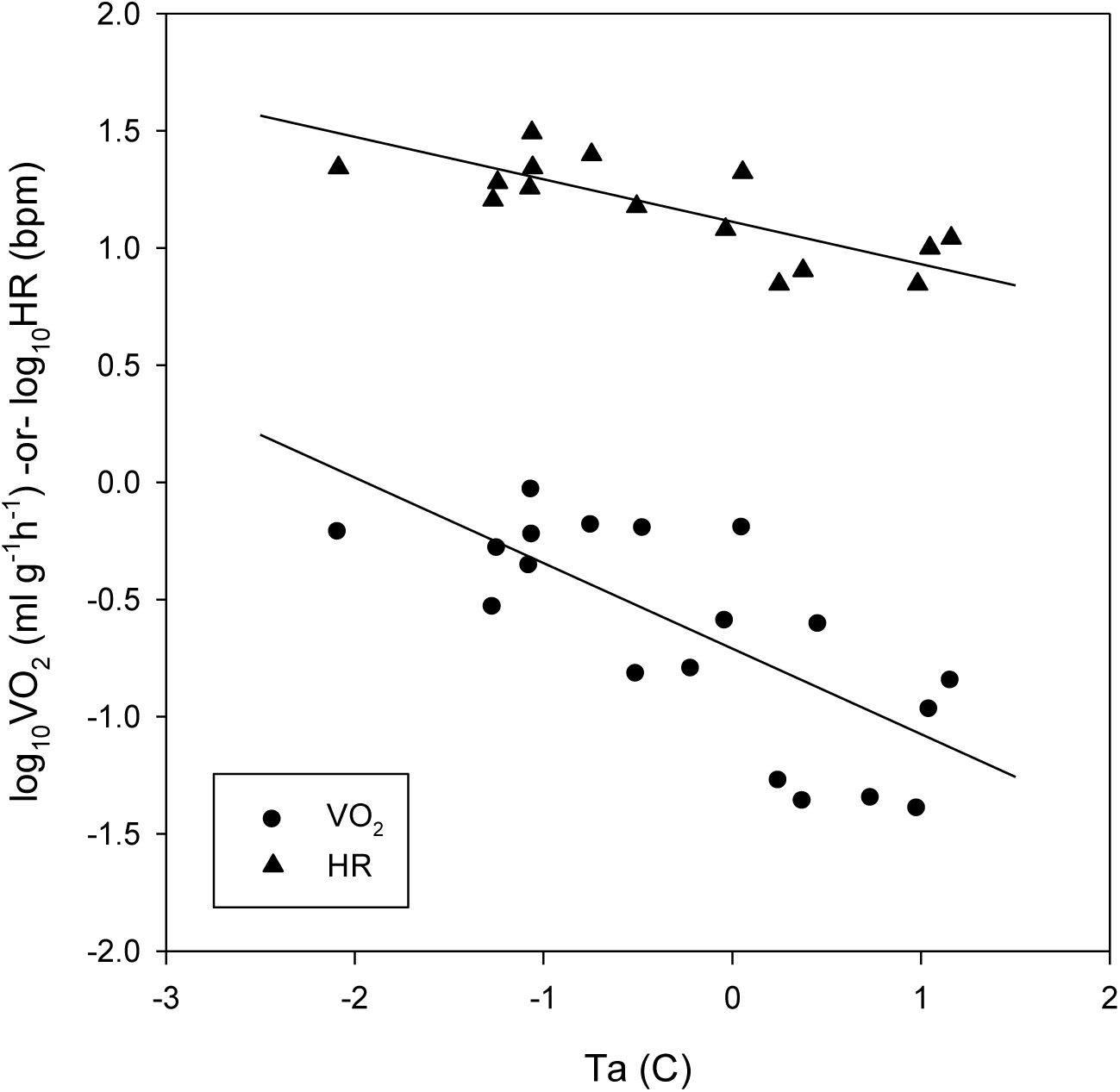
log transformed HR (triangles, dashed line; log_10_HR = 1.11 - 0.18(T_a_), r^2^ = 0.82, p<0.001) and 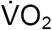 (circles, solid line; log_10_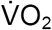 = −0.71 - 0.37(Ta), r^2^=0.71, p<0.001) for thermoregulating torpid bats as a function of T_a_. 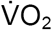 increased at a greater rate than HR below 1°C.

## Discussion

During thermoregulating torpor below T_crit_ animals not only require the ability to produce enough heat to maintain the possibly large gradient between T_b_ and T_a_, but also to ensure adequate cardiovascular function to support thermogenic needs. Our study is the first to provide simultaneously recorded data of metabolism and cardiac function in normothermic and torpid bats exposed to T_a_ at or below 0°C. Confirming our hypothesis, we found that torpid *C. gouldii* are capable of withstanding T_b_ down to at least 1°C while maintaining coordinated cardiac function. Below a T_a_ of ∼1°C animals began to defend T_b_ by increasing metabolic heat production and HR, and all animals in our study were capable of spontaneously rewarming from the lowest T_a_ (-2°C).

When animals were exposed to T_a_ below 1°C, there was a disproportionate response of the cardiovascular and metabolic systems as 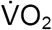 increased at a greater rate than HR to support elevated heat production. Bats in this study showed a substantial 46-fold increase in oxygen consumption, compared to a maximum increase of only 4-fold in HR. During this phase T_b_ remains low and the heart is cold. Although hibernators’ hearts are capable of withstanding very low T_b_, muscle contractility and rate are still limited by temperature (Michael & Menaker 1963; Smith & Katzung 1966; South & Jacobs 1973). In addition, it is possible that excessive cardiac stimulation and HR acceleration at low Tb may cause irritability of cardiac tissue and could possibly induce detrimental arrhythmias. Therefore, changes in circulation and the relationship between HR and 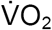 must be altered to maintain adequate supply under increasingly demanding conditions as T_a_ falls. In particular, the increased needs of the tissues must be met by an increase in oxygen delivery per heart beat or via accelerated oxygen extraction rates. Our results are the first to demonstrate a substantial upward shift in oxygen pulse during this phase of almost 15-fold, with HR only contributing to 31% of the elevated oxygen supply to tissues. Accordingly, we are also the first to illustrate a significant difference in relationship between HR and 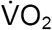 for thermoregulating torpid individuals compared to thermoconforming individuals with a greater 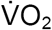 at almost every HR recorded for thermoregulating bats.

Our novel data highlight the differential responses of the cardiovascular and metabolic systems to thermogenic requirements at low Ta which would otherwise be unknown had we not recorded these variables simultaneously. Following the Fick equation (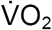 (ml/min) = HR × SV × (CaO_2_ - C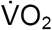)), the significant change in oxygen delivery per beat that we see in thermoregulating *C. gouldii* must be the result of substantial increases in both stroke volume (SV) and/or oxygen uptake by the tissues (arteriovenous difference;CaO_2_ - C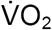). Due to the difficulties associated with measurements of stroke volume and blood characteristics, data for these variables in hibernating animals remain scarce with virtually no records available for bats in any physiological state. Under the assumption that SV increases to its maximum capacity (∼0.03 ml/min, calculated using Eq 7 of Bishop 1997; with the heart mass reported for *C. gouldii* from Bullen, McKenzie, Bullen & Williams 2009) in thermoregulating torpid bats we calculate that arteriovenous difference must increase to a maximum of 26.23 ml O_2_/dl blood at −2°C. This value is approaching maximal tissue oxygen extraction capacity, and is similar to oxygen extraction rates suggested during extreme exercise and flight (Bishop 1997). In order to achieve such high levels of oxygen extraction, haemoglobin (Hb) content of the blood must be at least 21 g/dl blood during torpor and 100% saturated in thermoregulating torpid bats. Previous reports of Hb content of the blood of bats vary greatly, with only one report for hibernating bats (Wołk & Bogdanowicz 1987), however values ≥20 g/dl blood have been reported in resting bats and would suggest our results are not outside of expectations for oxygen carrying capacity in these animals (Jurgens, Bartels & Bartels 1981; Arévalo, Pérez-Suárez & López-Luna 1987; Bishop 1997). In the wake of our findings for oxygen pulse, and in line with these calculations, our results suggest that both SV and tissue oxygen extraction approach maximal capacity when bats are thermoregulating during torpor at sub-zero Ta.

Additionally, the change in relationship between HR and 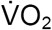 could also be the result of an alteration in blood supply across the body. During torpor, circulation to the periphery is restricted and perfusion of only vital organs such as the heart, lungs and brain is retained (Lyman & O'Brien 1963). This restriction of blood flow results in dramatic changes in blood pressure, enabling animals to maintain sufficient supply to tissues at low T_b_ when blood viscosity is increased (Lyman, Willis, Malan & Wang 1982). It is possible that when thermoregulating during torpor animals increase blood supply to essential thermogenic organs, such as brown adipose tissue or muscle, in order to defend Tb against increasing differentials with Ta. This may not only enable individuals to remain torpid for longer, but rewarm swiftly should the external temperature drop below a level they are capable of withstanding. It would be interesting to investigate how blood flow during this phase is altered, and whether the increased peripheral resistance and restriction of circulation is somewhat withdrawn. Our results show that in thermoregulating torpid bats thermal conductance is similar to that in resting bats, as was shown for *Myotis lucifigus* (Henshaw 1968) and this may be the result of such changes in blood flow. We have also previously suggested that this may be the case in bats during passive rewarming (Currie, Noy & Geiser 2015), and it may be a factor likely to influence the relationship between metabolism and heart rate under the conditions when animals are thermoregulating during torpor.

Our data are the first to show that *C. gouldii* readily express steady-state torpor and thermoconform below 10°C and we establish a new T_crit_ for thermoregulation of around 1°C, substantially lower than previous estimates for this species (Kulzer, Nelson, McKean & Möhres 1970; Hosken & Withers 1997). Consequently, we also present the lowest average 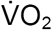 recorded for this species of 0.02 ml g^−1^h^−1^, which is only 20% of the previously recorded minimum (Hosken & Withers 1997). *C. gouldii* are capable of reducing HR to extremely low levels with an average thermoconforming minimum of 8bpm and absolute minimum of 3bpm recorded in one individual at T_a_ 1.0°C. This represents a 99.3% reduction in HR from the predicted basal HR for a 14g animal (422bpm) using the equation BHR = 816(W)^−0.25^ (Wang & Hudson 1971). These values are decidedly lower than anticipated for animals this size and compare to minimum torpid HR found in much larger hibernators (Lyman, 1951) *(Mesocricetus auratus* 110g, <15 bpm and *Ictidomys tridecemlineatus* 125g, 4-23 bpm; Lyman 1951) and marsupial hibernators at least twice their size *(Tachyglossus aculeatus* 4.8kg 7bpm; Augee & Ealey 1968; *Cercartetus nanus* 25-45g, 8bpm; Swoap, Körtner & Geiser 2017). Previously reported HR in torpid northern hemisphere bats ranged from 10-80bpm at 3-5°C (Reite & Davis 1966; Davis & Reite 1967; Kulzer 1967; Rauch 1973), while the lowest HRs previously reported were 4bpm in *Eptesicus fuscus* at 5°C (Rauch 1973; Twente & Twente 1978) and 5bpm in southern hemisphere *Nyctophilus gouldi* at 0°C (Currie *et al.* 2014).

When thermoregulating is activated during torpor, while still conserving substantial energy compared with normothermic thermoregulation at low T_a_ (99% less at −1°C), it is more energetically demanding than for thermoconforming torpid individuals (∼26-fold greater at - 1°C than 2°C). When both arctic-ground squirrels *(Urocitellus parryii)* and golden-mantled ground squirrels *(Callospermophilus lateralis* and *Callospemophilus saturatus)* were forced to thermoregulate during hibernation at increasingly lower T_a_ (down to −30°C), the frequency of spontaneous arousals increased and the amount of body mass lost during the hibernation season almost doubled compared to free-living animals (Geiser & Kenagy 1988; Richter *et al.* 2015). However, it has been suggested that the cost of arousals are comparatively reduced when animals are exposed to decreasing T_a_, as the costs of maintaining T_b_ during torpor increase (Karpovich *et al.* 2009; Richter *et al.* 2015). Conversely, unlike these rodents that hibernate in thermally stable hibernacula, *C. gouldii* roost in more thermally labile environments, such as tree roosts and buildings (Lumsden, Bennett & Silins 2002). All individuals from our study were originally removed from the roof of a building and were radio-tracked to tree roosts for at least 17 days following their release (Stawski & Currie 2016). Tree roosts and roofs of buildings are less insulated than caves or burrows and therefore are impacted by daily fluctuations in Ta, which can be dramatic on mild winter days (Law & Chidel 2007; Turbill 2008; Doty, Stawski, Currie & Geiser 2016). In these circumstances, bats are exposed to potential passive rewarming, thus mitigating much of the cost of arousal from low T_a_/T_b_. Indeed, we discovered evidence of passive rewarming in these same individuals when radio-tracked following their release from captivity, such that 83% of all recorded arousals during winter involved passive rewarming (Stawski & Currie 2016). Although poorly insulated roosts mean that animals may experience temperatures below their torpor Tcrit, the ability to passively rewarm may outweigh the costs of thermoregulation during torpor.

Previous investigations of HR and T_b_ in northern hemisphere bats exposed to low T_a_ have shown dramatic increases in HR below 0°C similar to the results of our study, however these studies do not present HR and MR simultaneously. Red bats *(Lasiurus borealis)* showed a similar response of HR to decreasing T_a_ as *C. gouldii* in this study, increasing from an average of 12 bpm at 5°C to 25-40 bpm at −2°C (Reite & Davis 1966). Similarly, data collected on metabolic rates in *L. borealis* during torpor at subzero Ta from a separate study are similar to our findings, at approximately 0.8 ml O_2_ g^−1^h^−1^ at −5°C (Dunbar & Tomasi 2006) compared to 0.61 ml O_2_ g^−1^ h^−1^ for *C. gouldii* at a T_a_ of −2°C. The physiological similarities we see between *L. borealis* and *C. gouldii* with regard to response to low T_a_ during torpor are likely reflective of their similar roosting habits over winter. *L. borealis* are also a tree-dwelling species and continue to remain in thermally labile roosts even though Ta may often fall below freezing (Mormann & Robbins 2007). These bats, like *C. gouldii*, also take advantage of mild winter days and are likely to passively rewarm (Dunbar & Tomasi 2006), but have been found to remain torpid unless external temperatures reach at least 15°C (Davis & Lidicker 1956; Davis 1970). On the contrary, little brown bats *(Myotis lucifugus)*, which overwinter in stable environments, were shown to respond to decreasing Ta by increasing HR 7-fold as Ta fell to −5°C (Reite & Davis 1966). At −2°C, HR was ∼100bpm in this species; up to 5-fold greater than bats in our study and the previously reported values for *L. borealis* (Reite & Davis 1966). However, these data may be more indicative of animals initiating or attempting the arousal process, as Reite and Davis (1966) note that after 4hrs animals kept at −5°C became hypothermic and died. This may suggest that bats which hibernate in unstable microclimates are more likely to remain torpid as Ta falls below zero, while bats that overwinter in caves are more likely to arouse as they can change roost location within the cave to avoid these excessively low temperatures.

Our data provide fundamental information regarding the costs of thermoregulating torpor, which has implications for understanding energy budgets of bats that regularly experience low T_a_ in the wild. Furthermore, disturbances such as sound (Speakman, Webb & Racey 1991), smoke (Stawski, Matthews, Körtner & Geiser 2015) and disease (Verant *et al.* 2014) have all been shown to increase T_set_ and therefore MR during torpor (Geiser 2004) as well as inducing arousal. As bats in the wild are increasingly susceptible to such disturbances it is crucial that we understand the costs associated with thermoregulating torpor as our data show that it is distinctly different from thermoconforming torpor.

## Conclusion

During the hibernation season animals may often be exposed to temperatures below their T_crit_. This is particularly true for tree-dwelling bats that roost in thermally labile environments. Importantly, our research provides valuable new information on MR and HR of hibernating bats over a range of temperatures and further advances our knowledge of the thermogenic capacity of small bats. Our results show that the T_crit_ of a temperate zone bat is most likely a response and selection to the long-term environmental temperatures of their habitat and can be close to zero, even though subzero temperatures may only be experienced for a few days throughout the year. Further, our data indicate that there is a significant difference in the physiological parameters of HR and 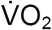 in thermoconforming torpid bats versus thermoregulating torpid bats. We also suggest that during thermoregulation in torpid animals tissue oxygen extraction and stroke volume must be altered in order to supply sufficient oxygen during this demanding phase, while the heart is rate limited by the cold and continues to beat relatively slowly.

## Acknowledgements

We would like to thank Heidi Kolkert for her help capturing bats, Anna Doty for her assistance with animal care and Brian Shaw for access to his property. We also appreciate the insights of Bill Milsom and Philip Currie on the study and their constructive comments on the manuscript.

## Funding

This research was supported by an Alexander von Humboldt Postdoctoral Research Fellowship awarded to SEC, an Australian Research Council Discovery Early Career Researcher Award to CS, and a grant from the Australian Research Council awarded to FG. The authors declare no conflicts of interest.

## Competing Interests

The authors declare no competing interests.

